# Genome-wide homology analysis reveals new insights into the origin of the wheat B genome

**DOI:** 10.1101/197640

**Authors:** Wei Zhang, Mingyi Zhang, Xianwen Zhu, Yaping Cao, Qing Sun, Guojia Ma, Shiaoman Chao, Changhui Yan, Steven S. Xu, Xiwen Cai

## Abstract

Wheat is a typical allopolyploid with three homoeologous subgenomes (A, B, and D). The ancestors of the subgenomes A and D had been identified, but not for the subgenome B. The goatgrass *Aegilops speltoides* (genome SS) has been controversially considered a candidate for the ancestor of the wheat B genome. However, the relationship of the *Ae. speltoides* S genome with the wheat B genome remains largely obscure, which has puzzled the wheat research community for nearly a century. In the present study, the genome-wide homology analysis identified perceptible homology between wheat chromosome 1B and *Ae. speltoides* chromosome 1S, but not between other chromosomes in the B and S genomes. An *Ae. speltoides*-originated segment spanning a genomic region of approximately 10.46 Mb was identified on the long arm of wheat chromosome 1B (1BL). The *Ae. speltoides*-originated segment on 1BL was found to co-evolve with the rest of the B genome in wheat species. Thereby, we conclude that *Ae. speltoides* had been involved in the origin of the wheat B genome, but should not be considered an exclusive ancestor of this genome. The wheat B genome might have a polyphyletic origin with multiple ancestors involved, including *Ae. speltoides*. These novel findings provide significant insights into the origin and evolution of the wheat B genome, and will facilitate polyploid genome studies in wheat and other plants as well.

## Introduction

Wheat (*Triticum aestivum*, 2n=6x=42, genome AABBDD), a major food grain source for humans, has been considered a typical allohexaploid originated from the interspecific hybridization involving three diploid ancestors (Sakamura 1918; Kihara 1919, 1954; Kihara et al. 1959; Sax 1922). This evolutionary theory of allopolyploid has led to successful identification of the ancestors for the wheat subgenomes A and D, but not yet for the subgenome B. The wild grasses *T. urartu* (2n=2x=14, genome AA) and *Aegilops tauschii* (2n=2x=14, genome DD) contributed A and D subgenomes to wheat, respectively (Kihara 1944; McFadden and Sears 1946; Dvorak et al. 1993). The ancestor of the subgenome B, however, remains controversial even though tremendous research efforts have been made to tackle this evolutionary puzzle of wheat for nearly a century.

Early studies on meiotic pairing, karyotyping, plant morphology, and geographic distribution of wheat-related wild species and their hybrids with wheat species (*Triticum* L.) identified the goatgrass *Ae. speltoides* (2n=2x=14, genome SS) as the closest ancestor of the wheat B genome (Jenkins 1929; Pathak 1940; Sarkar and Stebbins 1956; Riley et al. 1958). Meanwhile, questions had been raised about the meiotic pairing-based assessment of genome homology in these early studies because *Ae. speltoides* was suspected to contain genetic factors with epistatic effect on the wheat diploidization system, now designated *Ph* (pairing homoeologous) gene (Jenkins 1929; Sarkar and Stebbins 1956; Riley et al. 1958). The *Ph* gene limits meiotic pairing to homologous chromosomes in wheat and wheat hybrids with its relatives. Also, it had been assumed that the ancestral form of the B genome might have undergone a series of changes since its incorporation into wheat (Jenkins 1929; Sarkar and Stebbins 1956).

In a later study, Riley et al. (1961) confirmed the presence of the gene(s) in *Ae. speltoides* that suppresses the effect of the *Ph1* gene located on wheat chromosome 5B. Three genotypes with high, intermediate, and low ability to suppress *Ph1* activity were identified in *Ae. speltoides* (Dvorak 1972). According to these findings, Kimber and Athwal (1972) reassessed meiotic pairing in the hybrids and amphiploids involving wheat and *Ae. speltoides* accessions with different levels of suppression for the *Ph1* activity. They determined that the variation of meiotic pairing in the hybrids resulted from the presence of different *Ph1* suppressors in the *Ae. speltoides* accessions. Also, they found that chromosomes predominantly paired as bivalents in the amphiploid involving polyploid wheat and a low pairing *Ae. speltoides* accession, which was very similar to a normal diploidized allopolyploid. As a result, they concluded that *Ae. speltoides* could not be considered as the ancestor of the wheat B genome. This was also supported by the evidence of chromosome banding patterns (Gill and Kimber 1974) and protein electrophoretic profiles (Johnson 1972). More recently, three *Ph1* suppressor gene loci were identified and mapped to chromosome 3S, 5S, and 7S of *Ae. speltoides*, respectively (Dvorak et al. 2006).

In contrast, molecular analyses of both nuclear and extranuclear genomic DNAs suggested that *Ae. speltoides* or a species in the evolutionary lineage of *Ae. speltoides* could be the most likely ancestor of the wheat B genome as well as wheat plasmon (Ogihar and Tsunewaki 1988; Dvorak and Zhang 1990; Sasanuma et al. 1996; Wang et al. 1997; Kilian et al. 2007). However, comparative analysis of several gene loci and nearby genomic regions across the *Triticum* and *Aegilops* species did not reveal clear evidence supporting that conclusion (Hang et al. 2002; Salse et al. 2008). The *Ae. speltoides* S genome was considered to be evolutionarily closer to the wheat B genome than to the A and D genomes, but its candidacy as the ancestor of B genome remained undetermined in both studies.

The wheat B genome has significantly higher genetic variability than A and D genomes (Chao et al. 1989; Felsenburg et al. 1991; Siedler et al. 1994; Petersen et al. 2006). These findings support the hypothesis that the wheat B genome had diverged from its ancestor through various genomic modifications (Jenkins 1929; Sarkar and Stebbins 1956; Blake et al. 1999). In addition, *Ae. speltoides* has significantly higher intraspecific genetic variability than any of the other four *Aegilops* species in the Sitopsis section, which is even comparable to the interspecific variability among the other four *Aegilops* species in the section (Sasanuma et al. 1996). This appears to support the hypothesis that *Ae. speltoides* might contribute to the origin of the wheat B genome, but the current version of *Ae. speltoides* had diverged from the original ancestor of the B genome (Salse 2008). Another hypothesis, proposed by Zohary and Feldman (Zohary and Feldman 1962), states that the wheat B genome is a reconstructed genome resulted from meiotic homoeologous recombination between multiple ancestral genomes of the *Aegilops* species. This evolutionary recombination process was assumed to occur in the hybrids of the tetraploid amphiploids that combined the different ancestral *Aegilops* genomes and a common ancestral A genome of *T. urartu*. In other words, the wheat B genome might have a polyphyletic origin.

A species with a genome more closely related to the wheat B genome than the S genome of *Ae. speltoides* has not been discovered even though intensive search for the ancestor of B genome has been performed over nearly a century. It seems inevitable to reason that *Ae. speltoides* might contribute to the origin and evolution of the wheat B genome to some extent according to the previous studies. The present study aimed to assess the homology of individual wheat B-genome chromosomes with their homoeologous counterparts in the S genome of *Ae. speltoides* and to detect the *Ae. speltoides* genomic components in the wheat B genome if there are any. A novel integrative cytogenetic and genomic approach was taken to accomplish this research, which could not be done in the previous studies due to the lack of the genomics/cytogenetics tools and resources. This work shed new light on the origin and evolution of the wheat B genome, and will facilitate further studies of the complex polyploid genomes in wheat and other plants.

## Materials and Methods

### Plant Materials

Six “Chinese Spring” (CS) wheat B genome-*Ae. speltoides* disomic substitution lines [DS 1S(1B), DS 2S(2B), DS 4S(4B), DS 5S(5B), DS 6S(6B), and DS 7S(7B)] (Friebe et al. 2011), one substitution line involving chromosome 3S and 3A [DS 3S(3A)], and the CS *ph1b* mutant were the initial genetic stocks used in this research. They were provided by the Wheat Genetics Resource Center at Kansas State University, USA. DS 3S(3A) was included in this study because DS 3S(3B) is not available (please see results in this study). In addition, six CS wheat B genome-*Thinopyrum elongatum* (2n=2x=14, genome EE) disomic substitution lines [DS 1E(1B), DS 2E(2B), DS 3E(3B), DS 5E(5B), DS 6E(6B), and DS 7E(7B)] and one substitution line involving chromosome 4E and 4D [DS 4E(4D)] were used as controls to assess B-S genome homology in this study. DS 4E(4B) was not available. They were kindly supplied by J. Dvorak at UC Davis. Each of these disomic substitution lines has a pair of wheat chromosomes replaced by their homoeologous counterparts of *Ae. speltoides* or *Th. elongatum*. The common and durum wheat (*T. turgidum* ssp. *durum*, 2n=4x=28, genome AABB) accessions used in this study were selected from the worldwide diversity panel of the Triticeae Coordinated Agricultural Project (T-CAP). The other wheat species/accessions that contain the B genome were obtained from the U.S. National Plant Germplasm System. A total of 179 accessions under 13 wheat species (*Triticum* L.) were chosen based on their geographic origin and distribution, representing a diverse worldwide collection of the tetraploid and hexaploid wheat species (File S1). A subset of representative wheat species/accessions (n=88) were selected for single nucleotide polymorphism (SNP) genotyping from the 179 accessions (File S2).

### Construction of the special genotypes for meiotic pairing analysis

The CS-*Ae. speltoides* and CS-*Th. elongatum* disomic substitution lines were crossed and backcrossed with the CS *ph1b* mutant to construct the special genotypes monosomic for the individual B/A-S or B/D-E homoeologous pairs in the presence and absence of *Ph1*, respectively (Fig. S1). The chromosome-specific DNA markers (File S3) were employed to assist selection of the double monosomics for each of the homoeologous pairs. The selected individuals were verified for the monosomic condition by genomic *in situ* hybridization (GISH). The *Ph1*-specific DNA markers (Roberts *et al.*, 1999) were used to select the double monosomics with *Ph1* as well as those without *Ph1* (i.e. homozygous for *ph1b* deletion mutant) (Fig. S1).

Anthers with meiocytes [pollen mother cells (PMCs)] at metaphase I (MI) were collected for meiotic pairing analysis from the heterozygotes with and without *Ph1* following the procedure of Cai and Jones (1997). A total of over 100 meiocytes at MI from 1-6 plants were observed and analyzed for each of the special genotypes. GISH was used to differentially paint chromosomes of *Ae. speltoides*, *Th. elongatum*, and wheat for meiotic pairing analysis.

### DNA Marker Analysis

DNA samples were prepared as described by Niu et al. (2011). Chromosome-specific DNA markers, including SSRs (simple sequence repeats) and STSs (sequence-tagged sites), were developed and used for the identification of individual B-, S-, and E-genome chromosomes as described by Chen et al. (2007). Two STS markers (*PSR128* and *PSR574*) that tag the *Ph1* allele were used to identify individuals homozygous for the *ph1b* deletion mutant (Roberts et al., 1999). The wheat 90K iSelect SNP arrays were used to perform SNP genotyping assay for CS wheat, the disomic substitution lines, and the 88 representative wheat species/accessions (45 hexaploids and 43 tetraploids) (File S2) using the Illumina iScan instrument. SNP allele clustering and genotype calling were conducted using the GenomeStudio v2011.1 software (Illumina, Inc.) (Wang et al. 2014). The polymorphisms for each of the homoeologous pairs at the SNP loci were calculated as the percentages of the polymorphic loci out of the total loci genotyped. The graphical view of the genotypes for the 88 wheat species/accessions at the 68 SNP loci within the distal end of 1BL/1SL was constructed using the Flapjack software (Milne et al. 2010). Genetic diversity was calculated based on the SNP genotyping data of the 88 representative wheat species/accessions (Nei 1973) and plotted against the SNP consensus linkage map of wheat chromosome 1B (Wang et al., 2014). The SNP genotype-based cluster dendrogram was developed using R package “ape” (<u>https://cran.r-project.org/</u>).

### Fluorescent *in situ* hybridization (FISH)

Fluorescent genomic *in situ* hybridization (FGISH) was performed to differentiate wheat B-genome and S/E-genome chromatin from each other as described by Cai et al. (1998). Total genomic DNAs of *Ae. speltoides* and *Th. elongatum* were labeled with biotin-16-dUTP by nick translation as probe DNA for detecting *Ae. speltoides* and *Th. elongatum* chromatin, respectively. Total genomic DNA of CS wheat was used as blocking DNA. *Ae. speltoides*/*Th. elongatum* chromatin was painted with fluorescein isothiocyanate-conjugated avidin (FITC-avidin) as yellow-green and wheat chromatin was counter-stained with propidium iodide (PI) as red. Multicolor FISH was conducted following the procedure of Liu et al. (2006). The clone *pTa71*, a wheat 9 kb rDNA repeating unit that contains the 18S, 5.8S, and 26S rRNA genes and intergenic spacer (Gerlach and Bedbrook, 1979), was supplied by Peng Zhang at The University of Sydney, Australia. It was labeled with dig-11-dUTP and detected by anti-dig-rhodamine as red. This rDNA probe was used to tag the nucleolar organizer region on wheat chromosomes 1B and 6B. Total genomic DNA of *Ae. speltoides* was labeled with biotin-16-dUTP and detected by FITC-avidin as yellow-green. This genomic probe was used to identify *Ae. speltoides* chromatin in the wheat genome. Wheat chromatin was counter-stained with 4’,6-diamidino-2-phenylindole (DAPI) as blue. The fluorescence microscopy system BX51 (Olympus, Japan) was used to visualize GISH/FISH-painted chromosomes.

### DNA Sequence Analysis

The DNA sequences of wheat chromosome 1B was extracted from the IWGSC RefSeq v1.0 (<u>https://wheat-urgi.versailles.inra.fr/Seq-Repository/Assemblies</u>). The contextual sequences of the SNP loci at the distal ends of both CS wheat 1BL and *Ae. speltoides* 1SL were aligned to the DNA sequences of chromosome 1B using the Splign software (Kapustin et al. 2008). The physical order of the SNP loci was determined based on the DNA sequence alignment.

## Results

### Homology analysis of the individual B/A-S homoeologous chromosome pairs

Meiotic pairing has been considered direct cytological evidence for genome homology. It can, however, be influenced by the genetic factors in addition to homology, such as *Ph1* gene in wheat and *Ph1* suppressors in *Ae. speltoides*. To take account of the non-homology factors in the B-S genome homology analysis, we investigated meiotic pairing of individual B-S homoeologous pairs under the same genetic background of CS wheat in the presence and absence of *Ph1*. The CS wheat B genome-*Ae. speltoides* S genome disomic substitution lines dissect the S genome of *Ae. speltoides* into individual chromosomes in the CS wheat background. They were used to construct double monosomics for the individual B- and S-genome chromosomes. Meanwhile, the *ph1b* deletion mutant of *Ph1* was introduced into the double monosoimcs with assistance of the *Ph1*-specific DNA markers. Thus, we were able to investigate meiotic pairing of individual B-S homoeologous chromosome pairs in the presence as well as absence of *Ph1*. In addition, we investigated meiotic pairing of the individual B-E homoeologous chromosome pairs as controls for B-S genome homology analysis.

Chinese Spring wheat chromosome 1B was found to pair with *Ae. speltoides* chromosome 1S in 67 of the 134 PMCs analyzed (50.00%), while other B/A-S homoeologous pairs had a relatively low meiotic pairing frequency ranging from 0.00 to 8.63% in the presence of *Ph1* (Fig. 1). In addition, we noticed that 1B-1S meiotic pairing predominantly involved the long arms of chromosome 1B (1BL) and 1S (1SL). Surprisingly, wheat 1BL was found to contain a small *Ae. speltoides* S genome-derived chromosomal segment at its distal end, where meiotic pairing initiated (Fig. 2a). Also, the same *Ae. speltoides*-derived segment was observed on the unpaired 1BL (univalent) (Fig. 2b). The *Ph1* suppressor genes mapped to *Ae. speltoides* chromosomes 3S, 5S, and 7S (Dvorak et al. 2006). We found that meiotic pairing involving chromosome 5S was noticeably higher than that involving 2S, 3S, 4S, 6S, and 7S in the presence of *Ph1* (Fig. 1). Thus, there might be a *Ph1* suppressor on this particular *Ae. speltoides* chromosome 5S, but not on chromosomes 3S and 7S involved in this study. In the absence of *Ph1* (i.e. *ph1bph1b*), chromosomes 1B and 1S paired at a frequency of 60%, which was higher than the 1B-1S pairing frequency (50%) in the presence of *Ph1*. Meiotic pairing of other homoeologous pairs (2B-2S, 3A-3S, 4B-4S, 5B-5S, 6B-6S, and 7B-7S) was dramatically enhanced by *ph1b* mutant (Fig. 1). 5B^*ph1b*^-5S also exhibited a high pairing frequency (42.16%), suggesting absence of *Ph1* on *Ae. speltoides* chromosome 5S (Griffiths et al. 2006).

**Figure 1.**
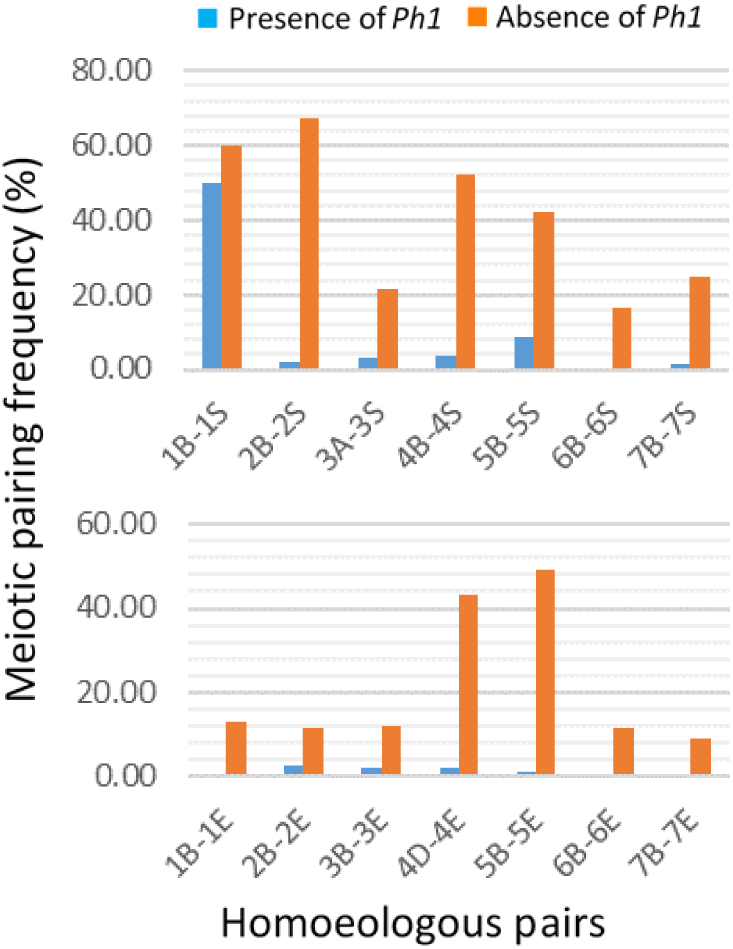
Meiotic pairing frequency of B/A-S (*top*) and B/D-E (*bottom*) homoeologous pairs in the presence and absence of *Ph1*.

Meiotic pairing was not observed between CS wheat chromosome 1B and *Th. elongatum* chromosome 1E in the 105 PMCs analyzed under the presence of *Ph1*. The other B/D-E homoeologous chromosome pairs also showed a low meiotic pairing frequency when *Ph1* was present (Fig. 1). Meiotic pairing of all B/D-E homoeologous chromosome pairs was enhanced by the *ph1b* mutant, but not as extensively as that with the B/A-S homoeologous chromosome pairs except 4D-4E and 5B-5E. Apparently, CS wheat chromosome 5B^*ph1b*^ had a high meiotic pairing affinity with *Ae. speltoides* chromosome 5S as well as *Th. elongatum* chromosome 5E (Fig. 1).

### GISH/FISH analysis of wheat B-genome chromosomes

Meiotic pairing analysis demonstrated a notable homology between CS wheat chromosome 1B and *Ae. speltoides* chromosome 1S (Fig. 1). Apparently, the *Ae. speltoides*-derived chromosomal segment at the distal end of 1BL contributed to the high 1B-1S pairing. To confirm the *Ae. speltoides* segment on 1BL and determine whether any additional *Ae. speltoides* segments are present on the CS B-genome chromosomes, we performed multicolor FISH/GISH to the mitotic chromosomes of CS wheat. The CS wheat chromosomes 1B and 6B were tagged by FISH using the rDNA probe *pTa71*; and *Ae. speltoides* chromatin was simultaneously painted by *Ae. speltoides* genomic DNA-probed GISH. An *Ae. speltoides* chromosomal segment was clearly detected at the distal end of CS 1BL, but not in other regions of chromosome 1B and other B-genome chromosomes (Fig. 2c). To further verify the origin of the distal segment on 1BL, we performed *Th. elongatum* genomic DNA-probed GISH to all three chromosome sets of the CS wheat genome. No *Th. elongatum*-derived GISH signals were observed on any of the CS wheat chromosomes, including 1BL (Fig. 3A). Thus, the distal segment on CS chromosomal arm 1BL is *Ae. speltoides* S genome-specific, not a common chromosomal region shared by CS wheat and its relatives. It was derived from the S genome of *Ae. speltoides*.

**Figure 2.**
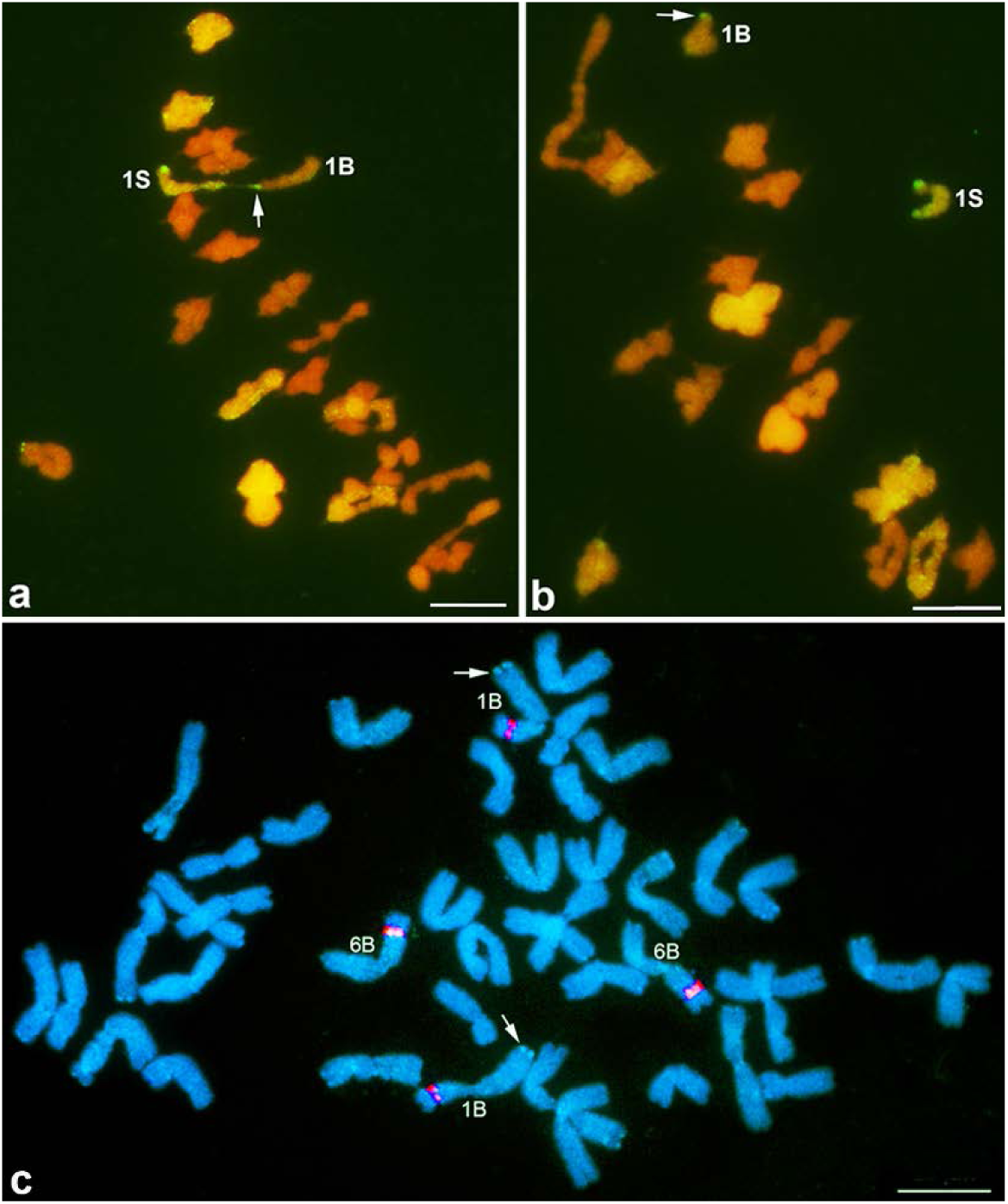
GISH/FISH-painted wheat and *Ae. speltoides* chromosomes. **a)** Showing meiotic pairing of CS wheat chromosome 1B with *Ae. speltoides* chromosome 1S as a rod bivalent; **b)** showing unpaired 1B and 1S chromosomes (univalents); and **c)** showing mitotic chromosomes of CS wheat. Arrows point to the *Ae. speltoides*-originated chromosomal segment on 1BL. Wheat and *Ae. speltoides* chromatin was painted as red and yellow-green by GISH, respectively (**a** & **b**). The *Ae. speltoides*-originated chromosomal segment was painted as light blue-green by GISH and the nucleolar organizer regions on 1BS and 6BS were painted as red by FISH. Wheat chromatin was counter-stained as blue by DAPI (**c**). Scale bar = 10 μm.

We surveyed the B genome of 179 representative accessions under 13 wheat species (*Triticum* L.) for the presence of *Ae. speltoides* chromatin by GISH. They were collected from the different geographic regions around the world and represented a diverse collection of the hexaploid and tetraploid wheat species/accessions that contain the B genome. All of these wheat species/accessions were found to contain an *Ae. speltoides* chromosomal segment at the distal end of 1BL as what we observed on 1BL of CS wheat, but not in the other regions of chromosome 1B and other B-genome chromosomes (Table 1; Fig. 2c). Therefore, the *Ae. speltoides*-derived chromosomal segment is universally present at the distal end of 1BL in both tetraploid and hexaploid wheat. It has been part of chromosome 1B probably since the incorporation of the B genome into tetraploid and hexaploid wheat.

**Table 1.** Presence of the *Ae. speltoides*-originated segment on chromosome 1BL in the 179 accessions under 13 wheat species (*Triticum* L.)

| Species | Genome | No. accessions | *Ae. speltooides* segment on 1BL* | Origin |
| --- | --- | --- | --- | --- |
| *T. aestivum* | AABBDD | 61 | + | Afghanistan, Algeria, Armenia, Australia, Belgium, Bhutan, Bosnia, Canada, China, Croatia, Czech Republic, Denmark, Egypt, Finland, France, Georgia, Germany, Greece, Hungary, India, Iran, Iraq, Israel, Italy, Japan, Kazakhstan, Macedonia, Mexico, Montenegro, Nepal, Pakistan, Russia, Serbia, Spain, Sweden, Switzerland, Syria, Tajikistan, Tunisia, Turkey, Ukraine, USA |
| *T. compactum* | AABBDD | 5 | + | Turkey, Syria, Egypt, USA |
| *T. macha* | AABBDD | 3 | + | Former Soviet Union, Iran, USA |
| *T. spelta* | AABBDD | 10 | + | USA, Ethiopia, Iran, Germany, Belgium, Afghanistan, Macedonia, Spain |
| *T. sphaerococcum* | AABBDD | 5 | + | Iraq, China, USA, Germany, Pakistan |
| *T. vavilovii* | AABBDD | 1 | + | Sweden |
| *T. dicoccoides* | AABB | 3 | + | Israel, Hungary |
| *T. polonicum* | AABB | 2 | + | Hungary, Jordan |
| *T. dicoccum* | AABB | 2 | + | Russia, USA |
| *T. carthlicum* | AABB | 2 | + | Iran, Georgia |
| *T. turgidum* | AABB | 2 | + | Ethiopia, China |
| *T. turanicum* | AABB | 2 | + | Morocco |
| *T. durum* | AABB | 81 | + | Argentina, Bulgaria, Mexico, Israel, Italy, Canada, Peru, Uzbekistan, Russia, Algeria, Azerbaijan, South Africa, Chile, Ukraine, Ecuador, Lebanon, United Kingdom, Ethiopia, France, China, India, Jordan, Malta, Oman, Hungary, Portugal, Pakistan, Saudi Arabia, Serbia, Nigeria, Morocco, Tunisia, Yemen, Turkey, Afghanistan, Portugal, Cyprus, Egypt, Eritrea, Iran, Iraq, Macedonia, North Africa, Pakistan |

\* “+” indicates presence of the *Ae. speltooides* segment on wheat chromosome 1BL.

### Comparative analysis of the individual B-S homoeologous pairs

Both CS wheat and CS-*Ae. speltoides* disomic substitution lines were genotyped using wheat 90K iSelect SNP arrays. The SNP genotyping results indicated that the substitution line originally designated DS 3S(3B) (Friebe et al. 2011) should be DS 3S(3A), which was further confirmed by SSR markers and chromosome C-banding. Thus, DS 3S(3A), instead of DS 3S(3B), was included in the SNP assay in addition to the substitution lines involving other six B-genome chromosomes (1B, 2B, 4B, 5B, 6B, and 7B).

High-throughput genotyping of CS wheat and the CS B genome-*Ae. speltoides* disomic substitution lines at 17,379 SNP loci identified a total of 6,722 SNPs polymorphic in the seven B/A-S homoeologous pairs. The homoeologous pair 2B-2S showed the lowest polymorphism (33.34%) and 7B-7S the highest (43.37%) at the SNP loci surveyed. The polymorphisms of other five B/A-S homoeologous pairs ranged from 33.89% (3A-3S) to 41.86% (4B-4S) (Fig. 4). Plotting of the SNP polymorphisms between CS wheat chromosome 1B and *Ae. speltoides* chromosome 1S against the SNP consensus linkage map of chromosome 1B (Wang et al. 2014) identified a genomic region that shared the same alleles at 65 of the 68 SNP loci within the distal ends of CS wheat 1BL and *Ae. speltoides* 1SL (Fig. 3C). Such a monomorphic linkage block was not detected in other chromosomal regions of the 1B-1S homoeologous pair and on other B/A-S homoeologous pairs (Fig. S2).

**Figure 3.**
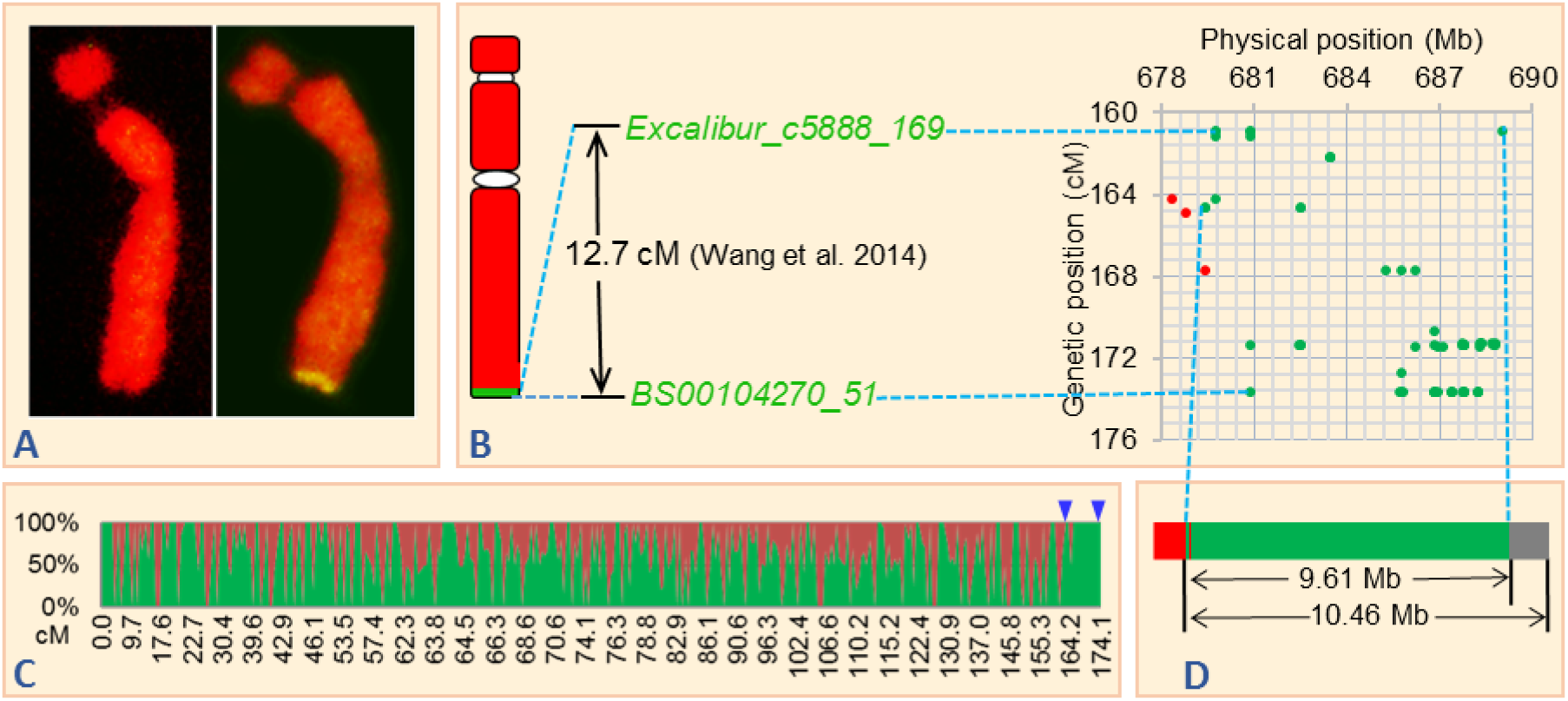
Cytogenetic and molecular mapping of the *Ae. speltoides*-originated chromosomal segment on 1BL. **A)** GISH-painted CS wheat chromosome 1B using *Th. elongatum* genomic DNA as probe (*left*) and *Ae. speltoides* genomic DNA as probe (*right*); wheat chromatin was painted as red and *Ae. speltoides*/*Th. elongatum* chromatin as yellow-green. **B)** Graphical representation of GISH-painted wheat chromosome 1B using *Ae. speltoides* genomic DNA as probe (*left*); genetic size of the highly monomorphic linkage block harboring 68 SNP loci at the distal ends of 1BL and 1SL (*middle*); and genetic and physical locations of the 68 SNP loci within the region (*right*). Green dots refer to the SNP loci monomorphic between CS wheat 1BL and *Ae. speltoides* 1SL; red dots refer to the polymorphic SNP loci. **C)** SNP-based comparative graph showing the distribution of polymorphisms between CS wheat chromosome 1B and *Ae. speltoides* chromosome 1S. Red areas refer to polymorphisms and green areas to monomorphisms. Arrow heads demarcate the highly monomorphic linkage block. **D)** Estimated physical size of the highly monomorphic linkage block harboring the 66 SNP loci and the extended region at the distal end of 1BL. Red, green, and grey bars refer to the polymorphic, monomorphic, and extended genomic regions, respectively.

**Figure 4.**
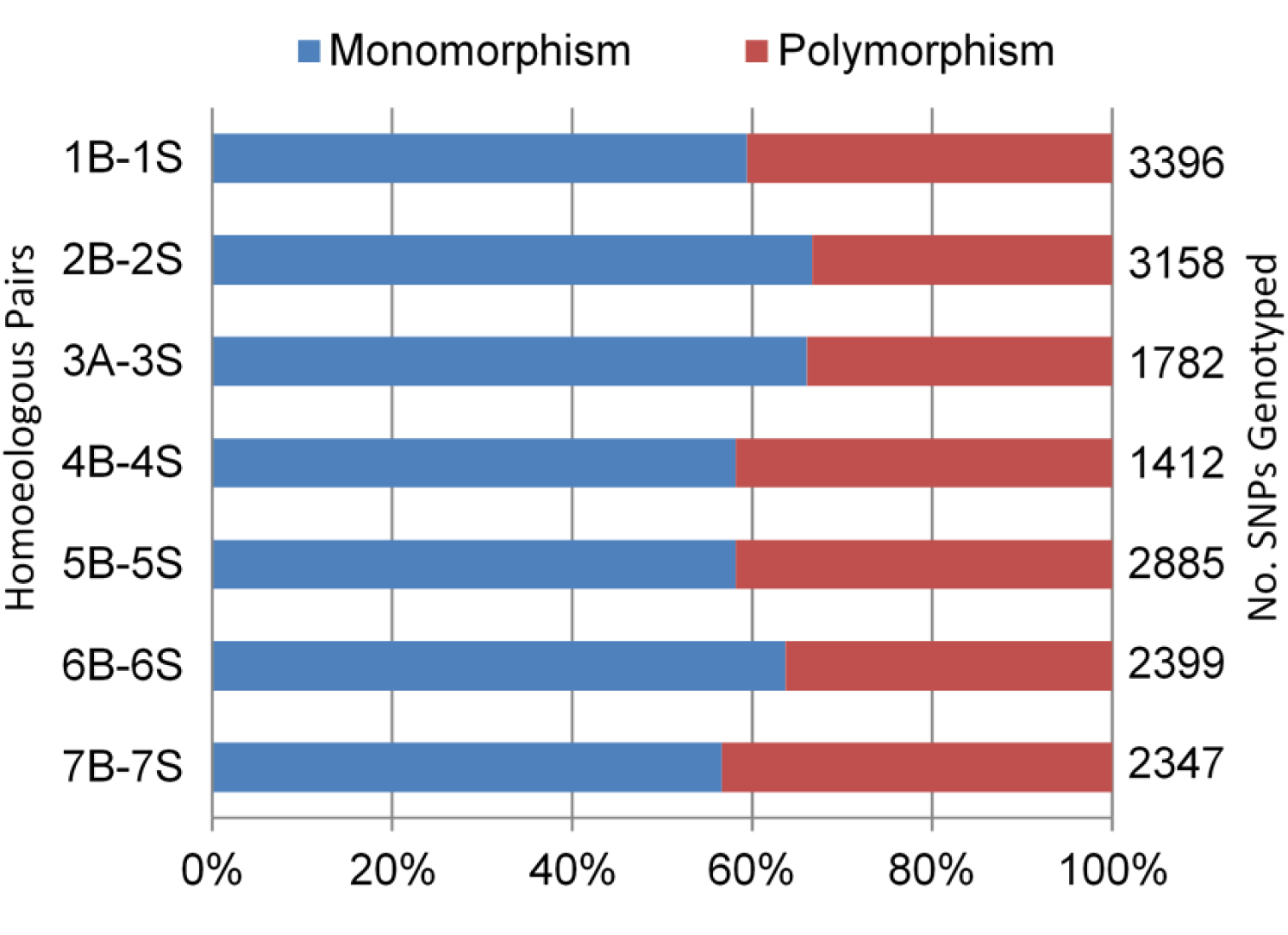
Polymorphisms of individual B/A-S homoeologous pairs at the SNP loci mapped on wheat B/A-genome chromosomes.

### Genetic and physical characterization of the 1BL distal end

The chromosomal regions at the distal ends of CS wheat 1BL and *Ae. speltoides* 1SL were found to be highly monomorphic at the 68 SNP loci (65 monomorphic/68 SNPs). The three polymorphic SNP loci within the region on 1BL and 1SL were positioned toward the proximal end of the region on the consensus linkage map (Wang et al. 2014). The entire region defined by the 68 SNPs spans a genetic distance of 12.7 cM (Fig. 3B and File S4). The contextual sequences of these 68 SNPs were aligned to the distal region of 1BL according to the IWGSC RefSeq v1.0 (<u>https://wheat-urgi.versailles.inra.fr/Seq-Repository/Assemblies</u>). Two of the three SNP loci polymorphic between 1BL and 1SL, *Tdurum_contig41999_2908* and *Ex_c1058_1537*, were physically assigned to the proximal end of the region. The other polymorphic SNP within the region, *RFL_Contig785_1156*, was distal to the first two polymorphic SNPs. One monomorphic SNP (*RFL_Contig785_1700*) was physically positioned within the interval between *Tdurum_contig41999_2908* and *RFL_Contig785_1156* according to the sequence alignment (Fig. 3B and File S4).

The genomic region that spans the 65 SNP loci monomorphic between CS 1BL and *Ae. speltoides* 1SL and one polymorphic SNP (*RFL_Contig785_1156*) was estimated to be 9.61 Mb in length according to the DNA sequence assemblies of 1BL (<u>https://wheat-urgi.versailles.inra.fr/Seq-Repository/Assemblies</u>). The two polymorphic SNP loci (*Tdurum_contig41999_2908* and *Ex_c1058_1537*) at the proximal end of that region were not included in the estimate (Fig. 3D). In addition, we identified an extended terminal segment of 0.85 Mb distal to the 68 SNP-defined region on 1BL according to the DNA sequence alignment. As a result, the total physical length of the SNP-defined and extended distal genomic region on 1BL was estimated to be 10.46 Mb (Fig. 3D). The actual physical size of this distal segment on 1BL might be greater than this estimate (10.46 Mb) because the current DNA sequence assemblies (<u>https://wheat-urgi.versailles.inra.fr/Seq-Repository/Assemblies</u>) we used in this study cover 689.9 Mb out of the total length 849 Mb of chromosome 1B (Šafář et al. 2010).

Four hexaploid wheat accessions (Cltr8347, PI429624, PI481728, and CI13113), similar to CS wheat, were found to be highly monomorphic with CS DS1S(1B) at the 68 SNP loci within the distal regions of 1BL and 1SL (Fig. S3). They were clustered together with CS wheat and CS DS1S(1B) in the dendrogram (Fig. 5). Cltr8347, PI429624, and PI481728 are the landraces from China, Nepal, and Bhutan, respectively. Both CS and Cltr8347 are the landraces collected probably in southwest China (P.D. Chen, personal communication), which is geographically close to the Himalayan region where Nepal and Bhutan are located. CI13113 is a winter wheat germplasm line with CS involved in the pedigree (<u>http://www.arsgrin.gov/npgs/</u>). Thus, these five hexaploid wheat accessions seem to share a similar origin of this particular *Ae. speltoides*-derived genomic region on 1BL. In addition, we found that 11 tetraploid wheat accessions share the same genotypes at the 68 SNP loci on 1BL (Fig. S3), making them clustered together in the dendrogram (Fig. 5). These tetraploid wheat accessions originated from two primary geographical regions, the Mediterranean Basin and South America (File S2). Overall, the tetraploid wheat accessions showed higher genetic variability than hexaploids in the 68 SNP-defined genomic region on 1BL according to the cluster dendrogram and genetic diversity analysis (Figs. 5 and 6).

**Figure 5.**
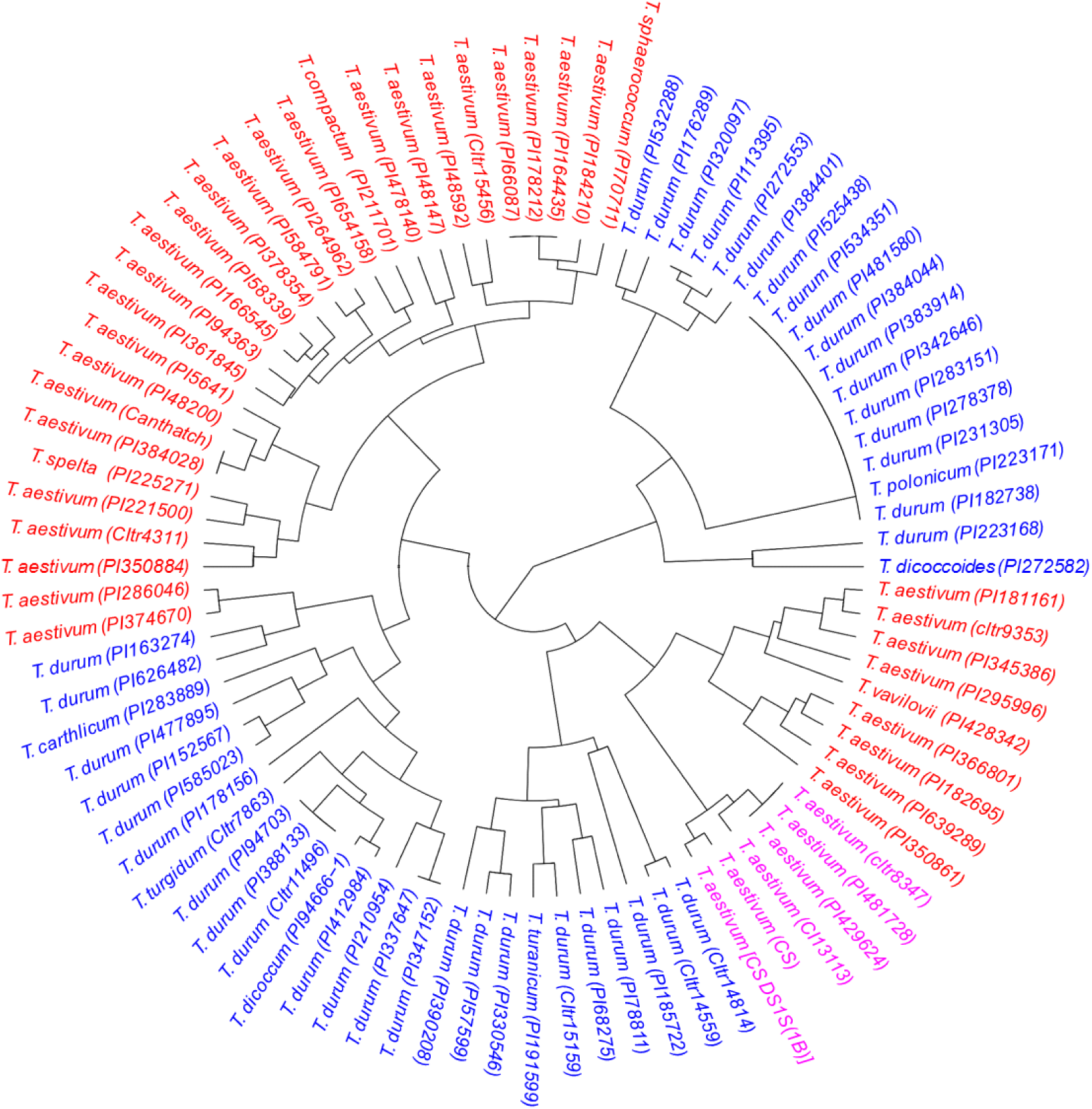
Cluster dendrogram of the 88 representative wheat species/accessions constructed based on the genotypes at the 68 SNP loci within the distal end of 1BL.

**Figure 6.**
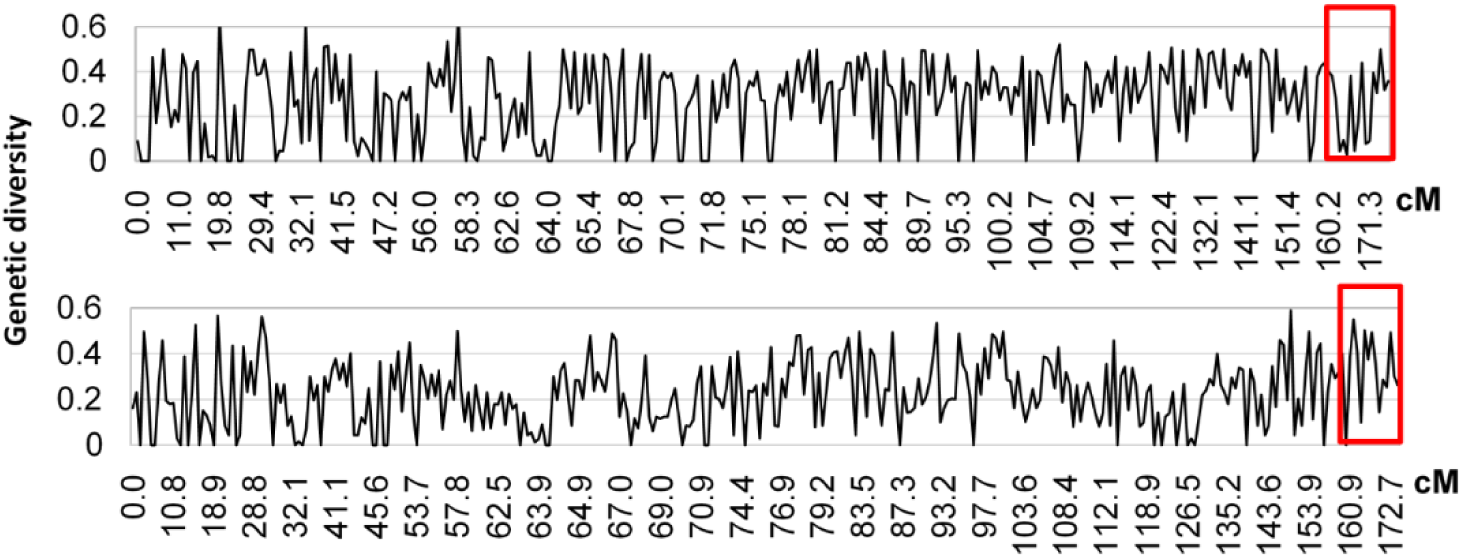
Genetic diversity at the SNP loci on chromosome 1B in 45 hexaploid wheat accessions (*top*) and 43 tetraploid wheat accessions (*bottom*). Red rectangles mark the *Ae. speltoides*-originated chromosomal region spanning the 68 SNP loci at the distal end of 1BL. Y-axis indicates genetic diversity defined as the probability of two different alleles randomly selected from the population.

Four of the 68 SNP loci (46805, 31066, 50867, and 76928) showed no allelic variation at all in the 88 wheat accessions. The other four SNPs (71971, 65270, 71898, and 78965) had very minimal variation in the wheat accessions (Fig. S3; File S4). Apparently, these SNP loci have been very conservative over the evolutionary process of this chromosomal region.

## Discussion

*Ae. speltoides* has been considered the diploid species with a genome most closely related to the wheat B genome according to the previous studies (Jenkins 1929; Pathak 1940; Sarkar and Stebbins 1956; Riley et al. 1958; Dvorak and Zhang 1990; Sasanuma et al. 1996; Wang et al. 1997; Kilian et al. 2007). However, the relationship of the *Ae. speltoides* S genome with the wheat B genome remains largely obscure. Previous meiotic pairing-based homology analyses had led to inconsistent conclusions about the evolutionary relationship of the B and S genomes due primarily to the influence of the *Ae. speltoides*-derived *Ph1* suppressors on meiotic pairing (Jenkins 1929; Riley et al. 1958; Kimber and Athwal 1972). In the present study, we partitioned the B and S genomes and investigated meiotic pairing of the individual B/A-S homoeologous pairs in the presence as well as absence of *Ph1*. This allowed us to monitor the effect of *Ph1* and *Ph1* suppressors in meiotic homoeologous pairing and to precisely assess homology for the individual B/A-S homoeologous pairs.

Substantial meiotic pairing was observed between CS wheat chromosome 1B and *Ae. speltoides* chromosome 1S in the presence of *Ph1* and absence of other S-genome chromosomes. The 1B-1S pairing frequency (50.00%) was significantly higher than any other B/A-S homoeologous pairs (0.00-8.63%). Chromosome 1S does not contain a *Ph1* suppressor gene (Dvorak et al. 2006). Therefore, no *Ph1* suppressor was involved in the 1B-1S meiotic pairing analysis. It was homology that made CS wheat chromosome 1B pair with *Ae. speltoides* chromosome 1S in a relatively high frequency. A small *Ae. speltoides*-derived chromosomal segment was detected at the distal end of CS wheat 1BL by GISH. Also, we found that 1B-1S meiotic pairing occurred mostly on the long arms (i.e. 1BL-1SL) that share the *Ae. speltoides* segment at the distal ends. Therefore, this *Ae. speltoides*-derived segment was the homologous counterpart on chromosomes 1B and 1S that initiated high meiotic pairing between them. Extremely low meiotic pairing (0 out of 105 PMCs) was observed between CS wheat chromosome 1B and *Th. elongatum* chromosome 1E in the presence of *Ph1*. Also, a *Th. elongatum*-specific segment was not detected on 1B and other B-genome chromosomes of CS wheat by GISH. As a control, these findings further confirmed the *Ae. speltoides* origin of the distal 1BL region in CS wheat.

We detected the *Ae. speltoides*-originated chromosomal segment at the distal end of 1BL in a representative worldwide collection of tetraploid (including wild and cultivated emmer wheat) and hexaploid wheat (n=179) in addition to CS wheat. Apparently, this *Ae. speltoides*-originated segment on 1BL has been part of the B genome in both tetraploid and hexaploid wheat species probably since the initial incorporation of the B genome into tetraploid wheat. Also, we found that tetraploid wheat had higher genetic diversity than hexaploid wheat within the *Ae. speltoides*-originated genomic region on 1BL, suggesting an evolutionary pattern similar to other genomic regions of polyploid wheat (Doebley et al. 2006). All these new findings consistently support the conclusion that *Ae. speltoides* had been involved in the origin of the wheat B genome. The *Ae. speltoides*-originated chromosomal segment on wheat chromosome 1BL has been retained in the B genome probably throughout the entire evolutionary and domestication process of polyploid wheat.

High-throughput SNP genotyping of individual B/A-S homoeologous pairs and the 88 representative wheat species/accessions identified a sizable highly monomorphic linkage block (12.7 cM) at the distal ends of wheat 1BL and *Ae. speltoides* 1SL. This was surprisingly consistent with the meiotic pairing and GISH results, supporting the *Ae. speltoides* origin of the distal segment on 1BL. The SNP-defined monomorphic genomic region and extended distal end on 1BL was estimated to be 10.46 Mb in physical size based on the genomic DNA sequence assemblies of chromosome 1B currently available from IWGSC RefSeq v1.0 (<u>https://wheat-urgi.versailles.inra.fr/Seq-Repository/Assemblies</u>). This estimate is about 1.2% of the total length of chromosome 1B (849 Mb) (Šafář et al. 2010). The GISH-detected *Ae. speltoides* segment on 1BL spans approximately 2.0% of the cytogenetic length of chromosome 1B. In addition, the DNA sequence assemblies of the IWGSC RefSeq v1.0 cover approximately 81.3% of the entire chromosome 1B (689.9 Mb out of 849 Mb) (Šafář et al. 2010). Thus, the monomorphic linkage block is probably a major portion of the *Ae. speltoides*-originated chromosomal segment on 1BL. The actual physical size of the *Ae. speltoides*-originated distal segment on 1BL might be greater than the estimate of 10.46 Mb.

Relatively low meiotic pairing was observed with each of other B/A-S homoeologous pairs in the presence of *Ph1*. In addition, we did not detect any *Ae. speltoides*-originated chromosomal segments in the other regions of chromosome 1B and on other B-genome chromosomes by GISH. Are there additional *Ae. speltoides*-originated chromosomal segments that are too small to leverage meiotic pairing and undetectable by GISH in the wheat B genome? We observed small monomorphic linkage blocks at multiple locations in other regions of chromosome 1B and on other B-genome chromosomes, but none was comparable in size to the one at the distal end of 1BL. Also, a clear association of those monomorphic linkage blocks with meiotic pairing could not be established in this study. Thus, we were unable to determine whether those B-genome chromosomal regions originated from *Ae. speltoides*. Further studies, such as genome-wide sequence comparative analysis, are needed to uncover the evolutionary relationship of those genomic regions with *Ae. speltoides*.

In summary, we conclude that *Ae. speltoides* had been involved in the origin and evolution of the wheat B genome. The current form of *Ae. speltoides* should not be considered an exclusive donor of the B genome. *Ae. speltoides* is probably one of the diploid ancestors involved in the evolutionary lineage of the B genome as stated in the theory of polyphyletic origin (Zohary and Feldman 1962). The wheat B genome might be a genome reconstructed from the homoeologous meiotic recombination between multiple ancestral genomes of *Aegilops* species, including *Ae. speltoides*. Further studies of *Aegilops* species, especially those in the Sitopsis section, may reveal additional insights into the origin and evolution of the wheat B genome.

## Author contributions

W.Z. and M.Z. equally contributed to this work on crossing, meiotic pairing analysis, SNP assays, GISH/FISH, comparative analysis of the homoeologous genomic regions, and manuscript preparation. X.Z. participated in crossing, chromosome-specific marker analysis, GISH, and manuscript preparation. Y.C. participated in GISH analysis. Q.S. performed computation in DNA sequence and comparative analysis. G.M. was involved in material maintenance and crossing. S.H. ran SNP assays and participated in manuscript preparation. C.Y. participated in DNA sequence and comparative analysis. S.S.X. was involved in data analysis and manuscript preparation. X.C. designed and coordinated this work, made crosses, analyzed and interpreted all experimental data, and prepared the manuscript.

## Acknowledgements

We thank members of the labs involved for their help to this research and Drs. Lili Qi and Rebekah Oliver for their critical review of the manuscript. This project is supported by Agriculture and Food Research Initiative Competitive Grant no. 2013-67013-21121 from the USDA National Institute of Food and Agriculture.

